# Cell-Cell-Seq resolves contact-associated NK cell activation in defined tumor cell dyads

**DOI:** 10.64898/2026.06.04.730259

**Authors:** Jesse Liang, Devon Shao, Dino Di Carlo, Joseph de Rutte

## Abstract

Cell-cell interactions shape immune recognition, but most single-cell transcriptomic methods measure cells after their interaction history has been lost or inferred. Here we apply Cell-Cell-Seq, a Nanovial-based workflow for sequencing defined cell pairs, to resolve contact-associated activation of natural killer cells paired with leukemia targets. Nanovials enabled controlled dyad formation, incubation, flow enrichment and droplet-based sequencing while reducing uncontrolled partner exchange and aggregation seen in suspension co-culture. Cell-Cell-Seq recovered a reproducible activation program marked by chemokine, cytokine, cytotoxic and immediate-early response genes. Compared with randomly mixed suspension co-culture, defined dyads emphasized contact-proximal activation, whereas suspension co-culture showed stronger features of early overstimulation. Dyad-resolved measurements also benchmarked computational models of cell-cell communication, identifying inferred signalling axes that were recovered and contact-induced programs missed by current approaches. These results establish Cell-Cell-Seq as a scalable strategy for mapping how defined immune-tumour encounters reshape cell state.

## Introduction

Emergent behaviors are a fundamental property of biological systems and have driven the development of technologies to advance our understanding of biology and ultimately develop enhanced therapeutics. Although the cell is widely regarded as the organizing unit of life, it is not until a cell interacts with another cell via molecular mediators such as cytokines, surface receptors and ligands, extracellular vesicles, and extracellular matrix that the defining behavior of tissues and organs characteristic of multicellular organisms emerges^1^. Understanding these interactions is particularly important in the context of disease, where dysregulation of cell-cell interaction is known to ultimately manifest in pathophysiology such as cancer and autoimmunity^2^.

A modern tool that has helped elucidate cell-cell interactions is single-cell RNA sequencing (scRNA-seq), which can capture the transduction and integration of cell-cell interactions at the mRNA and protein (CITE-seq^3^) levels. A paradigm shift enabled by single-cell measurements is the identification of rare and transitional populations that bulk measurements average away^4^, showing how disproportionately small subsets, like tumor-initiating cells, can drive tissue-level diseases such as cancer^5^. Despite the widespread proliferation of scRNA-seq techniques, critical challenges remain intrinsic to the technique for the purpose of interrogating emergent behavior. For example, the technology developed by 10x Genomics uses microfluidics to encapsulate cells, which have undergone some treatment or represent a patient-specific sample, into reaction droplets containing the necessary reagents to preserve the cell’s transcriptomic content^6^. The measurement of cells therefore loses spatial and neighbor context, which is vital to understanding the cell-cell interactions that underpin the emergent biology the technique is trying to elucidate.

Consequently, computational methods such as CellChat^7^, CellPhoneDB^8^, and LIANA/OmniPath^9^ arose to infer or prioritize ligand-receptor signaling potential from co-expression patterns. These approaches nominate candidate communication axes, but their outputs are difficult to validate because they are usually compared with other inferred interaction maps rather than a direct measurement of the transcriptional response produced by a defined cell pair. Tool-to-tool disagreement compounds this problem: the same dataset can return discordant interaction predictions depending on the inference framework applied^9^.

Spatial transcriptomics has emerged as a complementary technique^10,11^, retaining the context-rich environment of cells in their native tissue while providing single-cell resolution. However, preserving the entirety of the tissue also limits experimental control over interaction onset, partner number, and perturbation history. The resulting reference transcriptome is therefore the consequence of many hundreds, if not thousands, of cell-cell interactions across various time points, which adds a confounding element to the analysis and motivates computational efforts to infer causal or counterfactual tissue responses from spatial data^12^.

Large-scale single-cell atlases and perturbation datasets have also driven the development of cell foundation models trained to represent or reconstruct transcriptional states across cellular contexts^13^. Most current models are trained primarily on singlet measurements, including CRISPR Perturb-seq datasets^14^, and recent benchmarking studies have highlighted limitations in their predictive performance relative to established bioinformatic approaches^15,16^. One limitation of how these models are trained and evaluated is the relative absence of experimentally resolved cell-cell interaction states. Experimentally controlling the onset and duration of cell-cell contact offers a way to generate these missing measurements, linking interaction history to the transcriptional response that follows.

Including interaction-resolved datasets may improve the ability of these models to represent contact-dependent cellular responses. We therefore focused on individual interacting cell pairs (dyads) as the unit of analysis. Prior approaches such as PIC-seq demonstrated that physically interacting cell pairs can be isolated and sequenced to recover interaction-associated transcriptional states^17^. However, this approach, especially when isolating interacting pairs from tissues, is largely observational in nature and is precluded from controlling the onset of interaction and is confounded by other neighboring cells in the tissue of origin. To preserve physical cell pairing during sequencing in a workflow compatible with sorting and droplet-based scRNA-seq, we adapted Cell-Cell-Seq^18^, a dyad scRNA-seq workflow that uses Nanovials^19^ as hydrogel partitions to sustain cell-cell contact while permitting functional enrichment based on phenotypes such as target-cell killing and secretion.

The original work demonstrated these features in a PC3-T cell system and provided unique insights into the dynamic crosstalk between engineered effector and target cells, and featured computational strategies for deconvolving the shared transcriptome to individual cells. In our work we focused efforts on incorporated oligo-barcoded streptavidin tags for pooled-sample demultiplexing and higher-throughput dyad analysis. Further we chose a model system consisting of cytotoxic interactions between KHYG-1 natural killer cells^20^ and K562 leukemia cells^21^. This system is well suited for studying interaction heterogeneity because NK-mediated killing of K562 cells varies substantially across individual encounters despite the absence of obvious anti-killing ligands on the K562 surface^22^. By sequencing paired NK-K562 dyads, we measure contact-associated transcriptional responses and test how those measured responses relate to ligand-receptor inference and singlet-trained foundation-model reconstruction.

In this study, we report four connected results. First, we validate maintenance of paired-cell integrity through representative time-lapse imaging and flow-cytometry-based sorting measurements. Second, we apply the workflow to KHYG-1-K562 dyads and evaluate sequencing accuracy using concordance across Nanovial barcoding, Leiden clustering^23^, and CITE-seq surface-marker measurements, together with matched suspension co-culture controls. Third, we compare controlled Nanovial dyads with suspension co-culture to distinguish transcriptional programs associated with defined 1:1 contact from those associated with broader co-culture stimulation. Finally, we use dyad-resolved measurements as an experimental benchmark for ligand-receptor inference and singlet-trained foundation-model reconstruction, identifying both recovered communication axes and response programs that remain outside current computational representations.

Together, these results establish a workflow for profiling transcriptional programs in experimentally defined interacting cell pairs using commercially accessible single-cell technologies. More broadly, resolving how defined cellular encounters reshape transcriptional state could help characterize immune-tumor interactions and inform downstream applications such as target and biomarker discovery and therapeutic screening, while providing interaction-resolved data for models of cell-cell communication.

## Results

### Cell-Cell-Seq couples controlled Nanovial co-culture with dyad scRNA-seq

Cell-Cell-Seq was designed to measure the transcriptional response of defined cell pairs after a controlled period of co-culture. The workflow uses Nanovials as suspendable hydrogel partitions that can be loaded with two different cell types, incubated under conditions that preserve local, synchronized cell-cell contact, enriched by fluorescence-activated cell sorting, and processed by 10x Genomics single-cell sequencing with transcriptome and feature-barcode readouts (Figure 1). This design retains the core strengths of conventional scRNA-seq, including high-throughput cell-state measurement and compatibility with established droplet workflows, while adding an experimentally defined partner-history variable that is usually lost during library preparation. This compatibility builds on prior Nanovial studies demonstrating droplet-based single-cell RNA-sequencing and secretion-linked transcriptomic readouts^24,25,26^.

**Figure 1.**
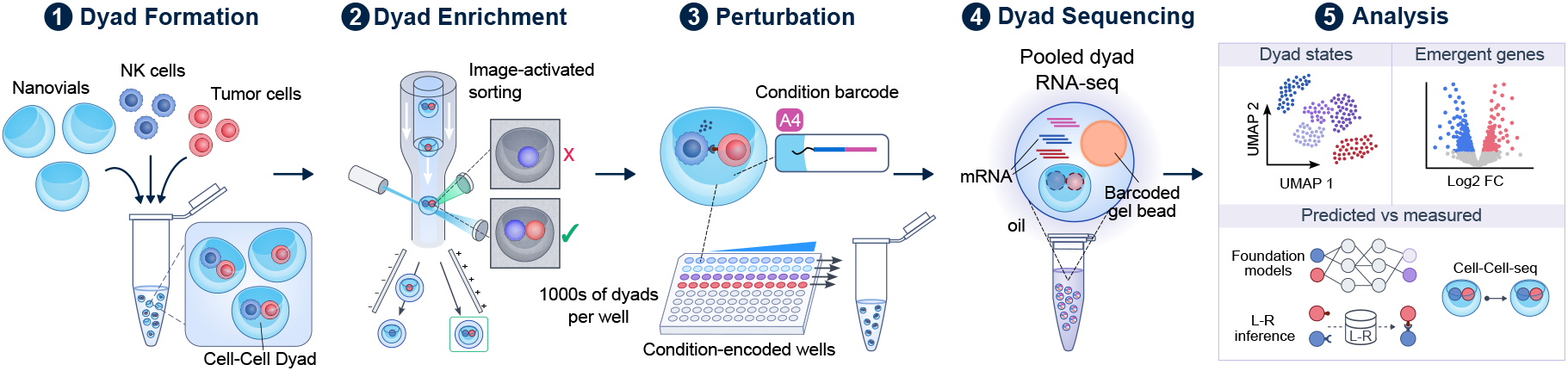
Cell-Cell-Seq workflow for profiling defined interacting dyads. Schematic overview of the Cell-Cell-Seq workflow. **1, Dyad formation:** NK cells and tumor cells are co-loaded into antibody-functionalized Nanovials to form local cell-cell dyads. **2, Dyad enrichment:** image-activated flow sorting enriches Nanovial events containing both cell types while excluding singly loaded or incorrectly loaded events. **3, Perturbation:** condition barcodes can be assigned upstream in plate-format workflows before pooling. **4, Dyad sequencing:** enriched dyads are processed by droplet-based single-cell RNA sequencing with transcriptome and feature-barcode readouts. **5, Analysis:** dyad-resolved measurements support cell-state assignment, identification of emergent contact-responsive genes, and comparison of measured dyad responses with ligand-receptor inference and foundation-model reconstruction.

We implemented this strategy in a KHYG-1-K562 immune-cancer interaction model. KHYG-1 cells provide a tractable natural killer (NK) effector population, while K562 cells provide a classical leukemia target that activates NK recognition in co-culture through reduced MHC-I (missing-self) recognition^27^. Matched suspension co-culture samples were included as comparators because bulk and flow cytometry-based NK-K562 activation assays are well described in the literature, but do not preserve the identity or number of target cells encountered by each NK cell. We hypothesized that transcriptomes generated from Nanovial and suspension co-culture would differ because suspension co-culture permits repeated target encounters, multicellular aggregation, and paracrine exposure histories that are not controlled at the single-cell level, with many interacting cells whose contacts are unsynchronized in time (Figure 1). The resulting experimental structure therefore allowed us to ask whether Nanovial dyad scRNA-seq could recapitulate the known KHYG-1-K562 activation response, and then whether controlling partner number revealed transcriptional structure that was obscured in conventional co-culture (Figure 1).

### Nanovials support recovery of intact interacting dyads while reducing uncontrolled partner exchange

To establish the physical basis for Cell-Cell-Seq, we first optimized Nanovial loading and recovery of KHYG-1-K562 dyads. KHYG-1 and K562 cells were captured on Nanovials using biotinylated antibodies against cell-type-specific surface markers. Notably, K562 loading was poor with beta-2 microglobulin and integrin-targeting antibodies that had previously supported efficient loading across several other human cell types, consistent with the low MHC-I presentation that contributes to K562 susceptibility to NK recognition^27^. After optimization, the final antibody combination (anti-CD71 to capture K562 and anti-B2M to capture KHYG-1) supported KHYG-1-K562 co-loading at upwards of 32% of Nanovial events as measured on a BD FACSDiscover S8 (Figure 2A) By comparison, spontaneous heterotypic dyads accounted for only ~0.1-0.7% of events in matched suspension co-culture (Figure 2B), one to two orders of magnitude lower than Nanovial co-loading..

**Figure 2.**
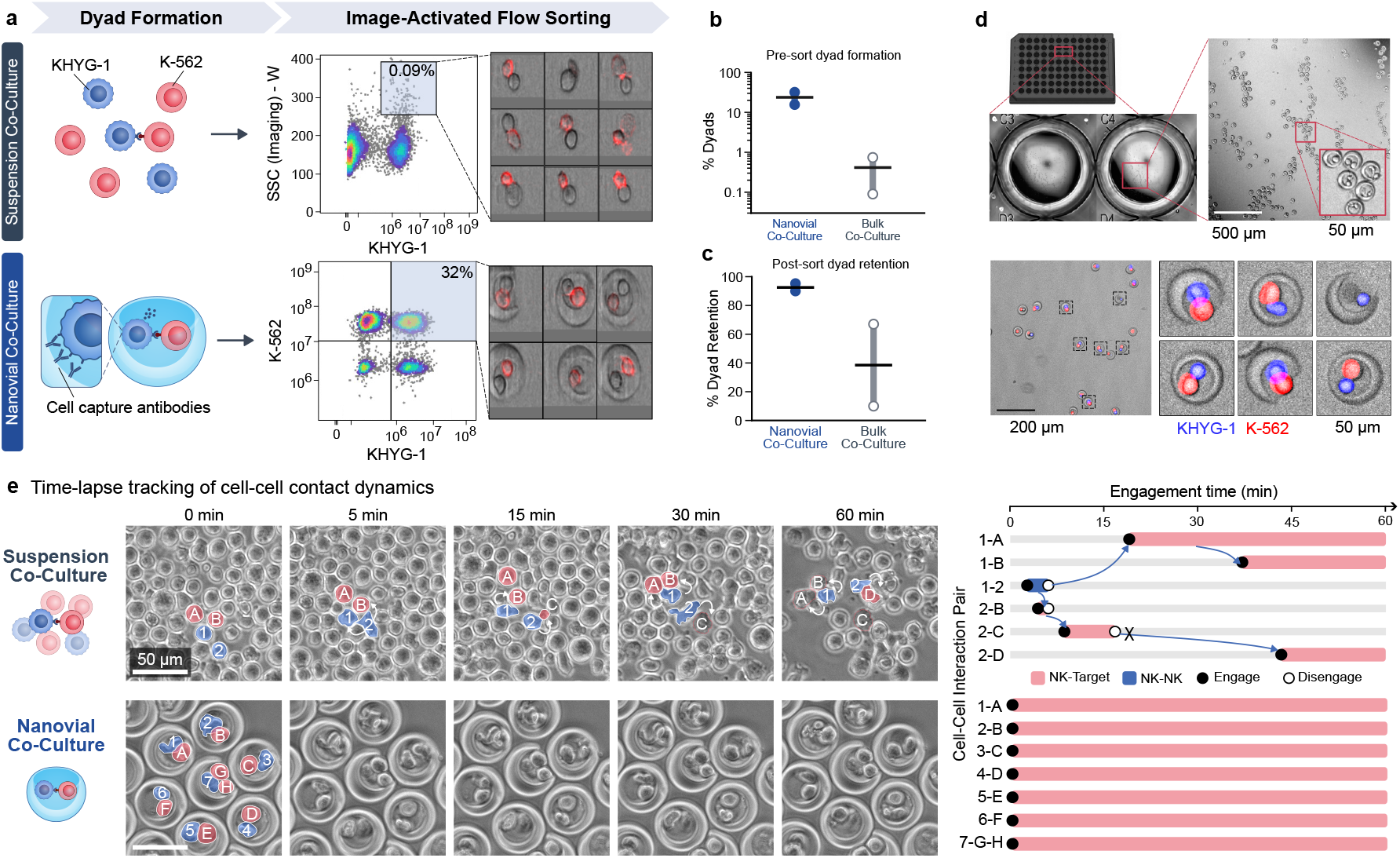
Nanovials enrich recoverable KHYG-1-K562 dyads and constrain partner exchange. **a**, Schematic and representative image-activated flow-sorting gates comparing spontaneous KHYG-1-K562 doublets in suspension co-culture with antibody-captured KHYG-1-K562 dyads in Nanovials. Representative images show the sorted event classes. **b**, Pre-sort dyad formation rates in Nanovial co-culture and bulk suspension co-culture. **c**, Post-sort dyad retention measured after recovery of sorted events. **d**, Representative microscopy images showing Nanovial-loaded dyads in microwells and after bulk recovery, with KHYG-1 and K562 fluorescence used to identify heterotypic pairs. **e**, Time-lapse imaging of suspension and Nanovial co-culture. Suspension co-culture shows partner exchange and repeated contact over the imaging window, whereas Nanovial co-culture maintains isolated cell pairs with constrained contact history.

Because heterotypic doublets can also occur spontaneously in suspension, a phenomenon other groups have exploited to study interacting immune cells by flow cytometry or single-cell sequencing^28,29,30^, we next compared Nanovial-associated dyads with heterotypic events observed in matched randomly mixed suspension co-culture conditions. Imaging on the BD FACSDiscover S8 confirmed that the sorting gates enriched for the intended heterotypic dyad events. We then quantified the proportion of sorted events that remained intact using widefield microscopy, demonstrating that KHYG-1-K562 pairs could be loaded, sorted, and recovered as Nanovial-associated dyads after flow enrichment (Figure 2B,C). Approximately 90-95% of sorted Nanovial events remained intact dyads, whereas retention of sorted suspension doublets was highly variable (~10-67%; Figure 2C), consistent with prior observations that such doublets frequently dissociate during sorting (~0.9% of sorted doublets remained intact post-sort in one report), a property that has itself been leveraged elsewhere to profile the separated partner cells^28,29^. Notably the higher rate of dyad events in Nanovials allows for higher throughput and bulk sorting of dyad-loaded Nanovials which allows for the design of 96-well plate scale experiments (Figure 2D).

We further interrogated the dynamics of each format using time-lapse imaging, which highlighted the partner-history difference between Nanovial and suspension co-culture. In suspension, individual NK cells moved between multiple K562 partners over the assay window, with manually traced examples contacting several target cells over time, consistent with prior observations that NK cells can serially engage and kill multiple targets in time-lapse assays^31^. In contrast, representative Nanovial time-lapse imaging supported constrained 1:1 physical pairing without the same partner exchange (Figure 2E). Notably, suspension co-cultures also produced large multicellular aggregates in several time-lapse experiments, whereas Nanovial-associated pairs remained physically isolated over the same imaging interval.

To test whether this physical-control workflow extended beyond the KHYG-1-K562 model, we also evaluated Nanovial pairing in a bispecific T-cell engager assay using Jurkat and Raji cells cultured with a blinatumomab biosimilar^32^ (Supplementary Figure 1 and Supplementary Video 6). As in the NK-K562 system, free-cell co-culture produced spontaneous aggregates visible by flow imaging and time-lapse microscopy, while the Nanovial workflow maintained isolated cell pairs during the assay window (Supplementary Videos 1–4). Together, these data establish the physical argument for Cell-Cell-Seq: Nanovials make interacting dyads recoverable before sequencing while reducing the uncontrolled partner exchange and aggregation that occur in suspension co-culture.

### KHYG-1-K562 dyad scRNA-seq recapitulates a reproducible NK activation program

We next asked whether sorted KHYG-1-K562 Nanovial dyads recovered an interpretable NK activation response. Sorted Nanovial events and matched co-culture samples were processed using 10x Genomics 5’ chemistry with feature-barcode libraries, allowing transcriptomes, sample tags, lineage markers, and surface-protein activation measurements to be analyzed from the same libraries (Figure 3A). For the Nanovial dyad experiments, oligo-barcoded streptavidin tags were used to pool matched 0-hour (dyad loaded Nanovials but kept on ice for the assay duration) and 2-hour co-culture dyad loaded Nanovial samples into a single 10x lane, reducing library-preparation batch effects while preserving sample identity. Transcriptome and feature-barcode libraries were generated for two independent Cell-Cell-Seq dyad experiments and a matched co-culture dataset, then processed using standard quality-control, normalization, clustering, and annotation workflows (Supplementary Figure 2, 3).

**Figure 3.**
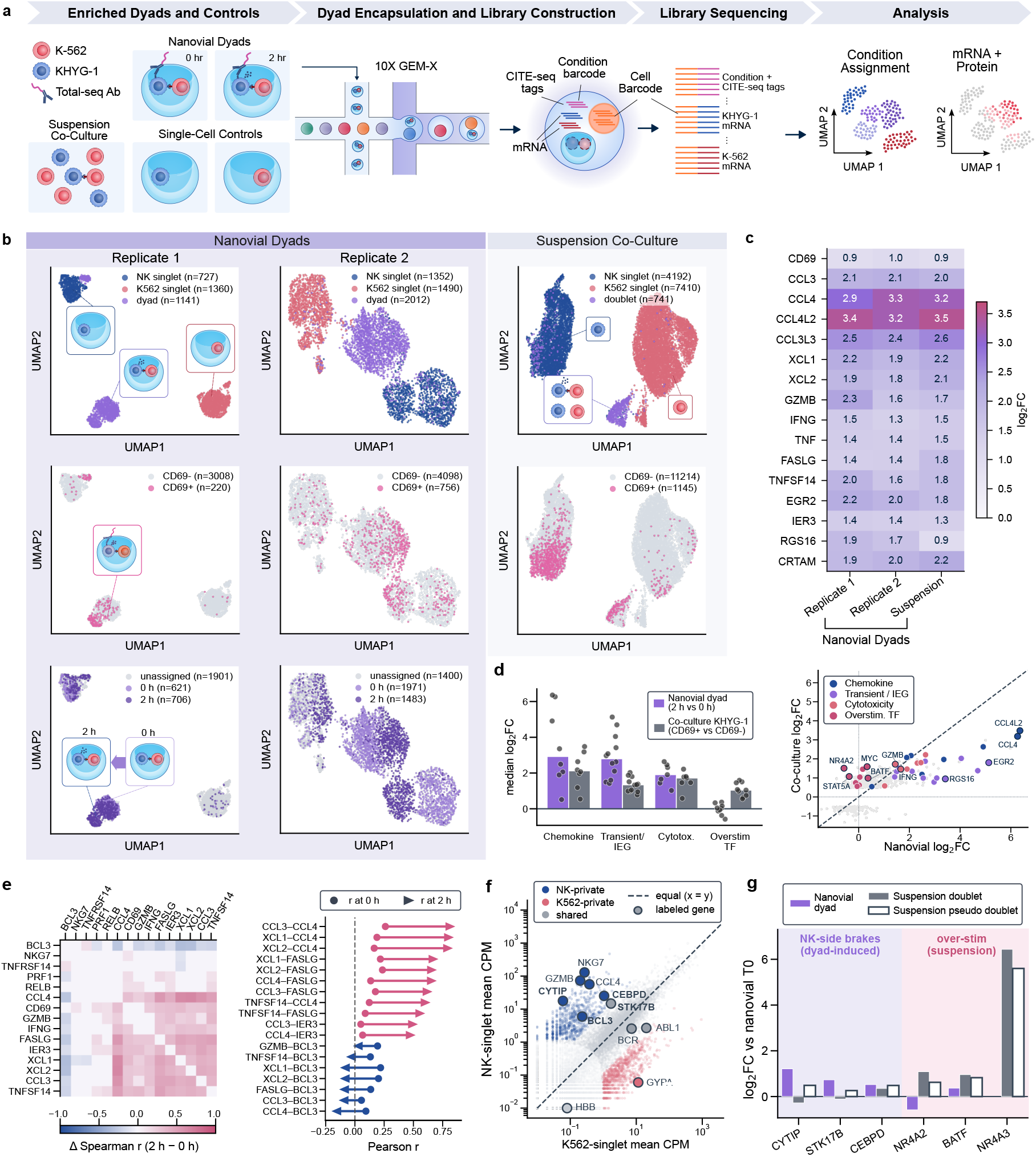
Cell-Cell-Seq recovers reproducible KHYG-1-K562 activation and distinguishes Nanovial dyads from suspension co-culture. **a**, Experimental and analysis workflow for dyad scRNA-seq. Sorted Nanovial dyads, matched single-cell controls, and suspension co-culture samples were processed with 10x transcriptome and feature-barcode libraries, enabling condition assignment, lineage annotation, and mRNA/protein analysis. **b**, UMAP embeddings for Cell-Cell-Seq replicate 1, Cell-Cell-Seq replicate 2, and suspension co-culture, showing assignment of NK singlets, K562 singlets, dyad/doublet-associated events, CD69 activation status, and Nanovial dyad time point. **c**, Heatmap of representative bactivation-associated genes across Nanovial dyad replicates and suspension co-culture. **d**, Program-level comparison of Nanovial dyad and suspension co-culture responses. Nanovial dyads show stronger median induction of chemokine, transient/immediate-early, and cytotoxicity programs, whereas suspension co-culture shows relatively greater induction of an overstimulation-associated transcription-factor program. **e**, Pairwise gene-correlation changes between 0 h and 2 h dyads, highlighting contact-associated rewiring among activation genes. **f**, Mean expression of genes in NK and K562 singlet populations, used to distinguish NK-private, K562-private, and shared transcript contributions in mixed dyad barcodes. **g**, Comparison of Nanovial dyads, suspension doublets, and pseudo-doublets generated from suspension singlets, showing dyad-induced NK-side programs and suspension-associated overstimulation markers.

Control Nanovials loaded with each individual cell type provided reference populations for sample demultiplexing and lineage annotation. These controls, together with oligo-streptavidin sample tags and literature-supported marker genes, enabled assignment of KHYG-1, K562, single-cell-loaded, and dyad-associated events in the pooled dataset (Supplementary Figure 4). UMAP visualization showed clear separation of the single-cell-loaded and dyad-associated populations, and CD69-positive events were enriched in the 2-hour co-culture condition (~32% of 2-hour dyads versus ~12% at 0 hours; Supplementary Figure 3D), consistent with contact-associated KHYG-1 activation (Figure 3B).

Activated KHYG-1 cells were defined using the reconciled CLR-normalized CD69 feature-barcode gate of >=1.2 across the reference dyad, sorted dyad, and co-culture datasets, providing a consistent protein-level activation threshold for cross-experiment comparisons (Supplementary Figure 3D)^33^. Across these samples, KHYG-1-K562 contact induced a conserved effector program. Differential expression recovered chemokines and cytokines associated with NK target recognition, including CCL3, CCL4, CCL4L2, CCL3L3, XCL1, XCL2, IFNG, and TNF; cytotoxic and death-ligand genes including GZMB and FASLG; and immediate-early response genes including EGR2, IER3, and RGS16 (Figure 3C)^34^. TNFSF14, encoding LIGHT, was also reproducibly induced^35^.

The activation response reproduced across Cell-Cell-Seq replicate 1 and Cell-Cell-Seq replicate 2. Among 167 genes called significant in both replicates, all 167 changed in the same direction, and activation effect sizes were strongly correlated (Pearson r = 0.78; Supplementary Figure 5). Gene-set enrichment analysis^36^ using the replicate 1 activation set against the ranked response in replicate 2 recovered the same program (NES approximately 2.9, permutation p < 0.001, leading edge 45 of 50 genes). The same canonical activation genes were also induced in co-cultured KHYG-1 cells, indicating that the activation signature reflects the KHYG-1-K562 interaction rather than a Nanovial-specific artifact. Thus, Cell-Cell-Seq preserves the expected NK activation biology while retaining information about the paired-cell context.

Pairwise gene-gene correlation analysis further showed that 2-hour dyads gained coordinated expression among activation-associated genes, supporting a program-level response within the dyad-associated state (Figure 3E). Because dyad-associated barcodes contain transcript contributions from both KHYG-1 and K562 cells, mean expression in annotated singlet populations was used to distinguish NK-private, K562-private, and shared genes, providing a reference for interpreting transcripts recovered from mixed dyad barcodes (Figure 3F). Using this framework, we compared Nanovial dyads with real suspension doublets and pseudo-dyads generated from suspension singlets as an additive-expression control. This analysis helped distinguish Nanovial dyad-induced NK-side responses from suspension-associated overstimulation signals (Figure 3G).

### Controlled dyads distinguish single-contact activation from co-culture overstimulation

Having established that Nanovial dyads and suspension co-culture shared a core KHYG-1 activation program, we next compared how each format distributed that response across contact-proximal activation and broader stimulation-associated transcriptional programs. In the Nanovial experiment, the primary response was defined as the log-fold change between 2-hour KHYG-1-K562 dyads and matched 0-hour dyads held on ice after loading. In the suspension co-culture experiment, the response was defined as the log-fold change between CD69-positive and CD69-negative KHYG-1 cells collected from the same 2-hour co-culture. These contrasts are not identical baselines, but they compare the transcriptional response to controlled 1:1 Nanovial contact with the activation state emerging from suspension co-culture, where KHYG-1 cells can repeatedly encounter multiple K562 targets and experience shared paracrine signals (Figure 2E)^31^.

Because Nanovial dyad barcodes contain RNA from both KHYG-1 and K562 cells, KHYG-1-private transcriptional responses are expected to be diluted relative to purified KHYG-1 singlets recovered from co-culture. We therefore interpreted cross-format comparisons conservatively, emphasizing within-format log-fold changes and program-level differences rather than absolute expression equivalence between dyads and purified KHYG-1 cells. This mixed-barcode structure should, if anything, bias against detecting larger KHYG-1-private responses in Nanovial dyads.

Using curated gene programs, Nanovial dyads showed larger median induction of chemokine genes (median logFC 3.36 versus 2.17 in co-culture), cytotoxicity genes (2.11 versus 1.75), and transient or immediate-early response genes (2.62 versus 0.99) (Figure 3D; Table 2). This dyad-enriched program included EGR2, RGS16, IER3, and CCL4, consistent with a synchronized contact-proximal response after defined KHYG-1-K562 pairing. In contrast, co-culture KHYG-1 cells showed relatively greater induction of the overstimulation-associated transcription-factor program (median logFC 0.99 versus 0.85 in Nanovial dyads), with NR4A2 and BATF preferentially induced in co-culture^37,38^. A separate enrichment analysis of system-specific genes also identified the overstimulation/AP-1-NR4A program among co-culture-enriched genes (odds ratio 36.46, p = 2.09e-5; hits: BATF, JUNB, NR4A2, NR4A3).

**Table 1.** Conserved KHYG-1 Activation Signature.

| Signature class | Genes | Interpretation |
| --- | --- | --- |
| Chemokines and lymphotactins | CCL3, CCL4, CCL4L2, CCL3L3, XCL1, XCL2 | Target-recognition-associated chemokine program induced in activated KHYG-1 cells |
| Cytokines | IFNG, TNF | Canonical NK effector cytokines induced after target-cell recognition |
| Cytotoxicity and death-ligand genes | GZMB, FASLG | Effector and target-cell killing-associated program |
| Immediate-early and contact-proximal response genes | EGR2, IER3, RGS16 | Early transcriptional response following KHYG-1-K562 contact |
| Surface/intercellular signaling genes | TNFSF14 | Reproducibly induced gene, retained as part of the activation signature but not interpreted as a mechanistic LIGHT-HVEM axis |

**Table 2.** Curated Program Definitions for Nanovial Dyad Versus Co-Culture Response Analysis.

| Program | Genes defined in analysis | Nanovial dyad median logFC | Co-culture KHYG-1 median logFC | Interpretation |
| --- | --- | --- | --- | --- |
| Chemokine | CCL3, CCL4, CCL4L2, CCL3L3, XCL1, XCL2, CCL5 | 3.36 | 2.17 | Larger in Nanovial dyads, consistent with a contact-proximal chemokine response |
| Transient/immediate-early | IER3, RGS16, EGR1, EGR2, EGR3, FOS, FOSB, JUN, JUNB, DUSP1, DUSP2, NR4A1, GADD45B, NAB2 | 2.62 | 0.99 | Larger in Nanovial dyads, consistent with synchronized early response capture |
| Cytotoxicity | GZMB, PRF1, FASLG, IFNG, TNF, TNFSF14, GNLY, NKG7, CRTAM | 2.11 | 1.75 | Shared activation program, modestly larger in Nanovial dyads |
| Overstimulation-associated TF | NR4A2, NR4A3, IRF4, BATF, PRDM1, ID2 | 0.85 | 0.99 | Relatively enriched in co-culture KHYG-1 cells |
| Checkpoint/exhaustion guardrails | PDCD1, HAVCR2, LAG3, TIGIT, KLRG1, CD96 | NA | NA | Used qualitatively to avoid overcalling terminal exhaustion at 2 h |

Single-cell marker detection provided a guardrail for this interpretation. NR4A3 was detected in 56% of co-culture CD69-positive KHYG-1 cells, compared with 18% of 2-hour Nanovial dyads and 1.6% of 0-hour dyads, while classical inhibitory receptors such as PDCD1, LAG3, and TIGIT remained low across groups. Thus, the co-culture-associated transcription-factor program is best interpreted as an early overstimulation-associated state rather than terminal exhaustion. Nanovial dyads therefore emphasize a synchronized 1:1 contact response, while co-culture KHYG-1 cells capture activation in a multicellular suspension environment where serial target encounter and paracrine exposure are less controlled.

### Dyad-resolved measurements benchmark interaction inference and singlet-trained foundation models

We next evaluated how dyad-resolved measurements relate to computational approaches that infer cell-cell interaction states from expression data and assessed whether single-cell foundation models trained primarily on singlet atlases capture cell-cell contact behavior (Figure 4A). We first applied LIANA to ask whether ligand-receptor inference from co-culture expression profiles could recover interaction programs measured directly by Cell-Cell-Seq. LIANA and related frameworks are useful for nominating candidate communication axes, but they infer signaling potential from co-expression and prior ligand-receptor knowledge rather than directly measuring the transcriptional consequence of a defined cell pair^9^. Cell-Cell-Seq therefore provides a controlled dyad-resolved benchmark for testing which inferred interaction axes are recovered, under-ranked, or absent from catalog-based annotations. We used KHYG-1 and K562 singlets from the suspension co-culture sample as the inference input, then compared the resulting ligand-receptor annotations with the contact-induced dyad response measured directly by Cell-Cell-Seq (Figure 4B-C).

**Figure 4.**
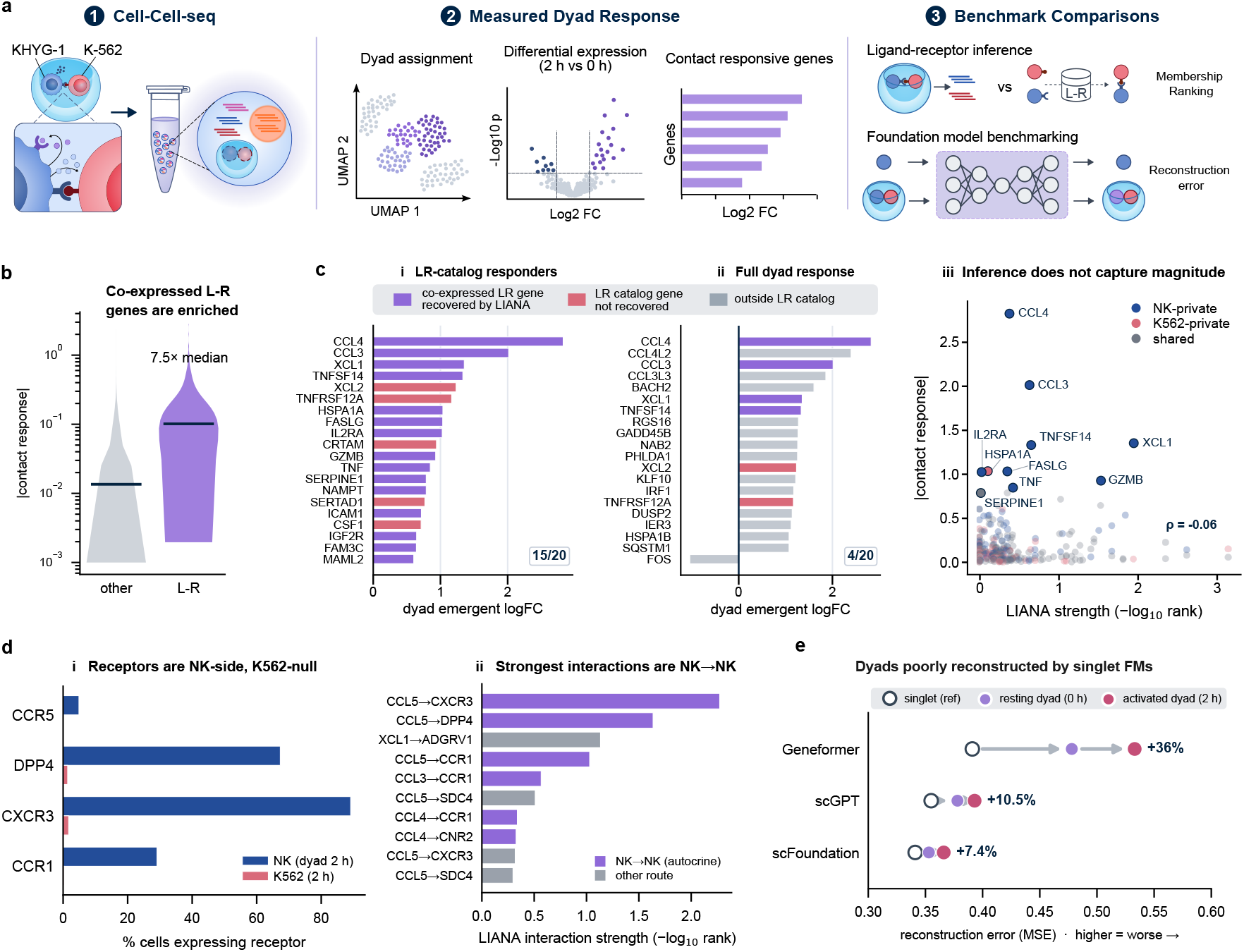
Dyad-resolved measurements benchmark ligand-receptor inference and singlet-trained foundation models. **a**, Schematic of the Figure 4 benchmarking strategy. Cell-Cell-Seq provides assigned KHYG-1-K562 dyads and measured contact-responsive gene programs, which are compared with ligand-receptor inference and foundation-model reconstruction. **b**, Co-expressed ligand-receptor genes inferred from suspension co-culture singlets are enriched among genes with measured Nanovial dyad responses. **c**, LIANA recovery within ligand/receptor catalog genes versus the full measured dyad response. LIANA recovers most of the strongest measured ligand/receptor-catalog responders, but LIANA-recovered ligand/receptor genes represent only a subset of the top contact-responsive genes overall, and LIANA interaction strength is not proportional to measured response magnitude. **d**, Receptor expression and inferred interaction direction support NK-to-NK/autocrine-like signaling potential among prominent chemokine axes, with receptors largely expressed on KHYG-1/NK cells and low or absent on K562 cells. **e**, Geneformer, scGPT, and scFoundation reconstruct activated dyads with higher error than held-out singlet references, indicating that contact-dependent dyad states are incompletely represented by singlet-trained foundation models.

This comparison showed that co-expressed ligand-receptor annotations were enriched among contact-responsive genes but did not explain the full dyad response. Using LIANA rank_aggregate inference with an expression-proportion threshold of 0.10, genes belonging to co-expressed ligand-receptor pairs had a 7.5-fold higher median absolute contact-induced log2 fold change than non-LR genes when LIANA was run on conventional suspension co-culture singlets (Mann-Whitney p = 1.86e-64; Fig. 4B). As expected, LIANA captured most of the strongest measured responders that were themselves ligand or receptor catalog genes, recovering 15 of the top 20 LR-catalog contact-responsive genes in the heterotypic LIANA result. However, this LR-restricted recovery did not extend to the full measured dyad program: LIANA-recovered ligand/receptor genes represented only 4 of the top 20 contact-responsive genes overall, and LIANA interaction-strength ranks were not proportional to measured response magnitude (Spearman rho = −0.061, p = 0.315; Fig. 4C). The recovered LR-catalog response included NK chemokines and TNF-family ligands, whereas many high-magnitude dyad-induced genes encoded intracellular transcriptional, stress-response, and feedback programs outside ligand-receptor catalog scope.

We next examined whether the residual ligand-receptor structure supported a heterotypic NK-to-K562 signal or an NK-intrinsic/autocrine-like program. Receptors for prominent chemokine axes were largely expressed on KHYG-1/NK cells and low or absent on K562 cells, and prominent inferred interactions included NK-to-NK/autocrine-like axes such as CCL5-CXCR3, CCL5-DPP4, XCL1-ADGRV1, and CCL5-CCR1 (Figure 4D). This pattern is consistent with NK-to-NK/autocrine-like signaling potential rather than a purely heterotypic NK-to-K562 interaction axis. Thus, Cell-Cell-Seq provides an experimental response layer that complements ligand-receptor inference: it distinguishes candidate signaling axes from realized contact-induced transcriptional outputs and can supply benchmark data for evaluating and improving communication-inference frameworks.

We also used the KHYG-1-K562 dataset as a test case for single-cell foundation models trained primarily on singlet atlases. Geneformer, scGPT, and scFoundation were evaluated using in-experiment singlets as a matched reference, followed by reconstruction analysis of 0-hour and activated Nanovial dyads (Figure 4E)^39,40,41^. Activated dyads reconstructed worse than held-out singlets across all three models, supporting the conclusion that pretrained singlet-atlas embeddings incompletely capture contact-dependent dyad states. The effect was largest for Geneformer, where median reconstruction error increased from 0.391 in singlets to 0.533 in activated dyads (+36%; Mann-Whitney p = 2.2e-69). The gap was smaller but still significant for scGPT (0.355 to 0.393; +10.5%; p = 3.2e-14) and scFoundation (0.341 to 0.366; +7.4%; p = 1.1e-7).

Together, these analyses show that Cell-Cell-Seq provides a direct benchmark for computational models of cell-cell interaction. Because Nanovial dyads define the interacting partners and contact interval before transcriptomic readout, inferred communication axes can be compared with the transcriptional programs that emerge during 1:1 contact. This pairing of prediction and measurement identifies both the ligand-receptor signals recovered by existing tools and the contact-induced response programs that remain outside current inference frameworks.

## Discussion

In this work we present Cell-Cell-Seq, a Nanovial-based dyad scRNA-seq workflow for measuring transcriptional responses from thousands of experimentally defined cell pairs. We used the Nanovial cavity to create a local co-culture environment in which paired cells could interact for defined intervals and then be enriched by flow cytometry. This controlled interaction window contrasts with suspension co-culture, where time-lapse microscopy showed repeated partner exchange and aggregate formation within the assay window in both KHYG-1-K562 and Jurkat-Raji pairing experiments. We then applied Cell-Cell-Seq to a KHYG-1-K562 immune-cancer interaction model and recovered a core NK activation program across two Cell-Cell-Seq replicates and matched suspension co-culture. However, the controlled dyad and suspension formats diverged in their stimulation-associated programs, with short 2-hour co-culture inducing a stronger NR4A/AP-1-associated overstimulation signature than matched Nanovial dyads. Finally, we used the same dyad-resolved measurements to evaluate ligand-receptor inference and singlet-trained foundation-model reconstruction, demonstrating how Cell-Cell-Seq can provide both a biological assay and a measured benchmark for computational models of cell-cell interaction. Together, these results motivate the themes we focus on below: how Cell-Cell-Seq extends paired-cell sequencing beyond proof-of-concept conjugate capture, how controlled dyad measurements separate biological interaction states that are otherwise collapsed in suspension co-culture, and how measured dyad responses can be used to evaluate computational representations of cell-cell contact.

### Scaling dyad-resolved sequencing through Nanovial partitioning

Nanovial partitioning makes dyad-resolved sequencing scalable by allowing defined cell pairs to be formed, incubated, enriched, and recovered before droplet-based transcriptomic readout. PIC-seq established that physically interacting cell pairs could be isolated and sequenced to recover interaction-associated transcriptional states^17^, providing an important foundation for paired-cell transcriptomics; however, throughput was constrained by the rate of native dyad formation and provided no means to control interaction time. Nanovials address this limitation by enabling controlled pairing and timed co-culture while remaining compatible with FACS enrichment and standard 10x feature-barcode workflows (Figures 1 and 2). This allowed paired-cell measurements to be collected at the scale of thousands of analyzed events in a format designed for routine droplet-based sequencing. Further increases in throughput and compatibility with 96-well plate assay formats would extend Cell-Cell-Seq toward perturbation workflows in which cell-pair identity, interaction time, and experimental condition can be varied across many parallel co-cultures.

### Defining interaction history reveals contact-resolved activation states

Suspension co-culture remains an important comparator because it is a standard format for NK cell killing assays and captures features of multicellular immune engagement, including repeated target encounter, serial killing, and paracrine exposure^31^. These same features can superimpose multiple interaction histories within a single transcriptional measurement, a challenge echoed by studies showing that paracrine signaling, microenvironmental control, and mixed stimulation reshape single-cell response states^42,43,44^. In the KHYG-1-K562 dataset, Nanovial dyads provided a complementary view by constraining partner number and coordinating the contact interval before sequencing. This timing likely helped resolve stronger induction of contact-proximal chemokine, cytotoxicity, and immediate-early response programs, including EGR2, RGS16, and IER3, whereas suspension co-culture showed relatively greater induction of an NR4A/AP-1-associated overstimulation program^37,38,45,46^ (Figure 3C,D). More broadly, Cell-Cell-Seq provides a path to disentangle the layered interaction histories that often arise in co-culture by measuring transcriptional responses after a defined cellular encounter.

### Dyad measurements as references for computational inference

Cell-Cell-Seq is complementary to approaches that preserve tissue context or infer communication from expression data, but its distinct value is that the interacting partners, contact interval, and partner-number history are experimentally defined before sequencing. Here, that structure allowed ligand-receptor inference to be evaluated against a measured dyad response rather than another inferred communication map. The result highlights both the utility and the boundary of catalog-based inference: LIANA recovered many ligand/receptor-class responders, while a substantial fraction of the contact-induced program consisted of non-ligand-receptor genes, including intracellular transcriptional, stress-response, feedback, and effector-response programs outside ligand-receptor catalog scope^9^ (Figure 4B,C). Extending this benchmark to frameworks that model broader communication networks, downstream target genes, or ligand activities, such as CellChat, NicheNet, CellPhoneDB, or related methods, could help determine which computationally nominated interaction axes are realized as transcriptional responses in defined cell pairs^7,8,47^. We also evaluated whether current singlet-trained foundation models reconstruct contact-dependent dyad states. Geneformer, scGPT, and scFoundation reconstructed activated dyads worse than held-out singlets (Figure 4E), suggesting that contact-dependent dyad states are not fully represented by models trained primarily on singlet atlases. These results underscore the need for collaboration between interaction-resolved experimental platforms and model developers, consistent with broader perturbation-modeling efforts showing that structured single-cell perturbation datasets are essential for training and evaluating response models^16,48^. Further work will be needed to determine the most useful format for paired-cell measurements in model training, whether as partner-cell context, paired-cell embeddings, interaction tokens, or other representations. Cell-Cell-Seq can contribute to that effort by generating datasets in which the interacting cells, contact interval, and transcriptional response are measured together.

### Scaling Cell-Cell-Seq across perturbations and cell states

Cell-Cell-Seq could also be extended to perturbation experiments. Because cell pairs are formed in Nanovials before sequencing, variables such as interaction time, effector identity, target identity, cytokine exposure, drug treatment, and sample barcode can be assigned before pooling for transcriptome and feature-barcode readout. This would allow parallel co-culture conditions to be compared while preserving a defined partner-history state for each condition, rather than relying on suspension cultures in which serial engagement and paracrine exposure vary across cells.

Longer term, Cell-Cell-Seq datasets could be used to train and evaluate models of contact-dependent immune states. Here, defined dyads were recovered, sequenced, and compared with suspension co-culture, ligand-receptor inference, and foundation-model reconstruction. Extending the workflow across primary donors, engineered immune cells, tumor backgrounds, time points, and perturbations could support applications in target validation, CAR-NK and T-cell engager development, cytokine and drug-response profiling, and selection of interaction programs that generalize across donor and disease contexts^49,50,51^. These datasets would also add interaction-resolved examples to single-cell atlases that are currently dominated by singlet measurements. Nanovial-based platforms have previously been applied to a range of functional single-cell and cell-interaction assays^18,52,53,54,55^. illustrating the breadth of cell-pairing and functional-enrichment approaches compatible with this platform.

Taken together, Cell-Cell-Seq makes partner history an experimentally accessible variable for single-cell analysis. Emergent biology arises when cells interact, yet most transcriptomic workflows measure cells after interaction history has been lost, inferred, or averaged across many encounters. By pairing Nanovial-based co-culture with flow enrichment and droplet-scale sequencing, Cell-Cell-Seq links defined cellular encounters to the transcriptional programs that follow. This provides a framework for mapping how cell-cell interactions shape multicellular behavior and for evaluating computational models against measured dyad responses.

## Methods

### Cell Line Sourcing And Maintenance

K562 (ATCC CCL-243), Raji (ATCC CCL-86), and Jurkat (ATCC TIB-152) cells were obtained from ATCC. KHYG-1 cells were obtained from Accegen Biotech (ABC-TC0506). K562 cells were maintained in IMDM, and KHYG-1, Raji, and Jurkat cells were maintained in RPMI 1640 (Invitrogen). Culture media were supplemented with 10% fetal bovine serum (Thermo Scientific) and 1x antibiotic-antimycotic (Santa Cruz Biotechnology). KHYG-1 medium was additionally supplemented with 100 U/mL recombinant human IL-2. Cells were maintained at 37 C with 5% CO2 between 1 x 10^5 and 1 x 10^6 cells/mL, with half-media changes to maintain log-phase growth for up to 25 passages. Frozen stocks were stored in vapor-phase liquid nitrogen, thawed, and passaged at least twice before use. Cell-line identity and mycoplasma status were based on vendor certification.

### Nanovial Modification

Partillion Bioscience 50 µm EZM Nanovials were used as cavity-containing hydrogel compartments for Nanovial co-culture experiments. Nanovial wash buffer, staining buffer (1% BSA in PBS), Nanovial modification, and cell loading were performed according to the Nanovial Developer Kit User Guide for Multicell Workflows (CF030) and SEC-seq-compatible Nanovial workflow principles^26^.

In brief, Nanovials were functionalized with streptavidin at a final concentration of 200 µg/mL, incubated for 30 minutes, washed three times by centrifugation, and coupled to biotinylated capture reagents selected for the cell-pairing system. KHYG-1-K562 experiments used α-B2M (Partillion Bioscience) for KHYG-1 and α-CD71 (Partillion Bioscience) for K562 loading, and Jurkat-Raji experiments used α-human integrin (Partillion Bioscience) for Jurkat and Raji loading. For experiments requiring Nanovial sample demultiplexing, oligo-barcoded streptavidin (BioLegend) was added to the streptavidin solution at a final concentration of 2.5 µg/mL.

### Cell Preparation And Cell Loading Into Nanovials

Cells were labeled before or during loading to enable cell-type discrimination by flow cytometry, imaging cytometry, and microscopy. For Jurkat-Raji bispecific experiments, Jurkat cells were stained with Calcein-AM and Raji cells were stained with CellTracker Red-Orange according to the manufacturers’ instructions at final concentrations of 1 µM in serum-free media for 15 minutes at 37 C.

For Cell-Cell-Seq replicate 1, KHYG-1 cells were stained with CellTracker Deep Red at 1 µM for 15 minutes in serum-free media at 37 C, and K562 cells were stained with Calcein-AM at 1 µM for 15 minutes in serum-free media at 37 C. KHYG-1-only and K562-only Nanovial controls were generated using the same loading procedure.

For Cell-Cell-Seq replicate 2, K562 cells were stained with CellTrace Yellow (Invitrogen) at a 1:500 dilution of stock dye for 20 minutes at 37 C. KHYG-1 cells were labeled after Nanovial loading with HLA-B-RB705 antibody (BD Biosciences; 5 µL antibody in 100 µL staining volume) for 15 minutes on ice.

Cells were co-loaded into Nanovials at a 1:1:1 Cell A:Nanovial:Cell B ratio according to the Partillion Bioscience Developer Kit User Guide (CF030). In brief, Cell A was added at 1 x 10^6 cells/mL to the Nanovial suspension, mixed by pipetting 10 times, centrifuged at 100 x g for 1 minute, and incubated on ice for 15 minutes. The suspension was mixed again by pipetting 10 times, centrifuged at 100 x g for 1 minute, and incubated on ice for an additional 15 minutes. Cell B was then added, mixed by pipetting 10 times, centrifuged at 100 x g for 1 minute, and incubated on ice for 15 minutes. The full suspension was incubated for an additional 15 minutes to complete loading.

### Flow Cytometry And Fluorescence-Activated Cell Sorting

Nanovial and suspension co-culture samples were analyzed and sorted on a BD FACSDiscover S8 using a 130 µm nozzle. Nanovial event identification followed Section 4 of the Partillion Bioscience Demonstrated Protocol for Adjusting Cell Ratios to Optimize Nanovial Multicell Loading (CF031). In brief, Nanovial-associated events were separated from non-Nanovial events using SSC (Violet)-A and Lightloss (Violet)-A, followed by gating on the large Nanovial population. Double-loaded Nanovials were identified using the fluorescence channels corresponding to each cell type, and image-enabled cytometry was used to verify that sorted gates contained the intended heterotypic Nanovial-associated events. Sorting followed established strategies for isolating Nanovials on commercial flow sorters^56^.

For suspension co-culture dyad identification, SSC (Imaging)-W was used to identify doublet events containing KHYG-1 signal. This gate was used because KHYG-1 homotypic doublets were rarely observed under the tested conditions. Target events, defined as dyad-loaded Nanovials or suspension dyads, were sorted in purity mode into either 15 mL conical tubes containing PBS with 0.04% BSA for 10x Genomics library generation or into media-containing 96-well plates for imaging.

### Microscopy And Time-Lapse Imaging

Post-sort dyad retention was quantified by manually counting dyad and singlet events after sorting into 96-well plates. Phase-contrast images were acquired on an EVOS M5000 microscope (Invitrogen) using a 10x objective.

Wide-field images of sorted dyad-containing Nanovials were acquired on a Cytation 5 using a 4x PL FL objective. A 3 x 3 montage was collected per well, stitched with the native Cytation 5 linear-blend tool, and exported for analysis in ImageJ.

Time-lapse imaging was performed on a Cytation 5 (BioTek) using a 10x PL FL phase objective. For prolonged imaging and culture, media were supplemented with 25 mM HEPES and the environmental chamber was maintained at 37 C. For bispecific-driven workflow-validation experiments, Jurkat and Raji cells were co-cultured with a blinatumomab biosimilar (100 ng/mL; ProteoGenix) and imaged for 7 hours at 8 minutes 20 seconds intervals. For KHYG-1-K562 experiments, cells were cultured in a 50:50 mixture of the corresponding culture media and imaged for 90 minutes at 15 seconds intervals.

Time-lapse image compilation and analysis were performed in ImageJ. Individual cells were manually labeled and tracked over the indicated imaging intervals to quantify the number of interactions and partners encountered by individual KHYG-1 cells. KHYG-1 cells were assigned by morphology and behavior relative to K562 cells, which typically appeared spherical and comparatively static.

### Co-Culture Conditions And Single-Cell Gene-Expression And Feature-Barcode Library Generation

For Cell-Cell-Seq replicate 1, four Nanovial samples were prepared and encoded with oligo-barcoded streptavidin. After loading, one dyad-loaded Nanovial sample was incubated for 2 hours at 37 C with 5% CO2. During this incubation, KHYG-1-only Nanovials, K562-only Nanovials, and 0-hour dyad-loaded Nanovials were maintained on ice and sorted as described above for downstream library preparation. After the incubation period, 2-hour dyad-loaded Nanovials were sorted for dyad-associated events. 0-hour and 2-hour dyad-loaded Nanovial populations were then stained with oligo-barcoded anti-CD69 according to the manufacturer’s recommendations. All samples were pooled into a single 10x Genomics 5’ GEM-X lane, followed by GEM generation on a Chromium iX and library preparation according to the Chromium GEM-X Single Cell 5’ Reagent Kits v3 with Feature Barcode technology for Cell Surface Protein user guide (CG000734). Libraries were sequenced on a NovaSeq instrument by MedGenome.

For Cell-Cell Seq replicate 2, Nanovial samples included the same conditions as well as a suspension co-culture sample consisting of KHYG-1 and K562 cells mixed at a 1:1 ratio. Unlike replicate 1, in replicate 2 loaded dyads were sorted first and then incubated at physiological conditions for two hours. Following the incubation period, all samples were stained with oligo-barcoded α-CD69 and taken through the same workflow specified above, except that three lanes of the 10x were used: (1) single-cell loaded controls, (2) dyad-loaded Nanovials at the 0-hour and 2-hour time points, and (3) suspension co-culture at 2 hours.

### Preprocessing And Cell Annotation

Raw sequencing files were processed using the Cell Ranger pipeline through the 10x Genomics Cloud Analysis platform to generate gene-expression and feature-barcode count matrices. These matrices were then analyzed in Python using the scverse ecosystem. Transcriptome, sample-tag, lineage-marker, and activation-marker information were imported at the same barcode level, allowing each recovered barcode to be annotated using both RNA and feature-barcode measurements.

Cells with low transcript recovery were removed, using a minimum threshold of 3,000 detected genes for the sorted-dyad library to account for the higher RNA content expected from dyad-associated barcodes. Genes detected in fewer than three cells were excluded. Gene-expression counts were normalized, log-transformed, and used for dimensionality reduction, clustering, and UMAP visualization.

Cell identities were assigned by integrating oligo-barcoded sample tags, lineage-associated transcriptome scores, and feature-barcode measurements. KHYG-1 and K562 lineage scores were calculated from canonical marker genes, and score distributions were thresholded with a two-component Gaussian-mixture model. Barcodes with high KHYG-1 and K562 scores were classified as dyad-associated events, while single-lineage barcodes were classified as KHYG-1 or K562 singlets (Supplementary Figure 2). Timepoint identity was assigned from the SAv-C0971 and SAv-C0972 Nanovial hashtags (BioLegend), corresponding to 0-hour and 2-hour samples, respectively. Barcodes were assigned to a timepoint when the dominant hashtag had at least 50 UMIs and was at least threefold higher than the next highest hashtag; remaining barcodes were classified as hashtag doublets or negatives. CD69 activation status was assigned using a CLR-normalized CD69 feature-barcode threshold of 1.2, corresponding to the Q88 threshold of the 0-hour-dyad null and applied consistently across datasets.

### Differential Expression And Signature Scoring

Differential expression was used to define response programs within each experimental format. For Nanovial dyads, the response was defined by comparing 2-hour dyads with matched 0-hour dyads. For suspension co-culture, the KHYG-1 response was defined by comparing CD69-positive and CD69-negative KHYG-1 cells within the 2-hour co-culture sample. Differential expression was performed using Wilcoxon rank-sum testing with Benjamini-Hochberg correction. Genes were required to be detected in at least 10% of cells in the relevant comparison to reduce fold-change inflation from sparsely detected transcripts. For downstream binary gene-set summaries, significant up-regulation was defined as adjusted p < 0.05 and log2 fold-change > 0.5.

Replicate conservation was evaluated by comparing activation-associated response genes between Cell-Cell-Seq replicate 1 and Cell-Cell-Seq replicate 2. In each replicate, CD69-positive and CD69-negative dyad-associated events were defined using the matched 0-hour-dyad CD69 null distribution, and response-gene direction and effect size were compared across replicates.

Signature scores were calculated from curated gene modules defined from the differential-expression results and prior literature. Module scores were compared across 0-hour Nanovial dyads, 2-hour Nanovial dyads, suspension co-culture doublets, and CD69-positive KHYG-1 cells from suspension co-culture. Checkpoint and exhaustion-associated genes were evaluated as a separate marker set from the overstimulation-associated transcription-factor module.

### Ligand-Receptor Inference And Benchmarking

Ligand-receptor inference was performed using LIANA with the OmniPath/consensus ligand-receptor resource and rank-aggregate scoring. KHYG-1 and K562 singlets from the suspension co-culture sample were used as the primary inference input. Ligand-receptor pairs were retained when the ligand and receptor each passed a minimum expression-proportion threshold of 0.10 in the corresponding sender and receiver populations.

LIANA outputs were benchmarked against the measured Cell-Cell-Seq dyad response, defined as differential expression in 2-hour Nanovial dyads relative to 0-hour Nanovial dyads. For gene-level comparisons, retained ligand-receptor complexes were decomposed into component genes, and genes belonging to co-expressed ligand-receptor pairs were compared with all other genes by absolute measured contact-response magnitude. LIANA interaction strength was also compared with measured dyad-response magnitude to evaluate whether ligand-receptor prioritization was proportional to the experimentally observed response.

To distinguish recovery within the ligand-receptor catalog from recovery of the full dyad response, measured contact-responsive genes were evaluated in two scopes. First, contact-responsive genes that were members of the ligand-receptor catalog were assessed for recovery by LIANA. Second, the full set of contact-responsive genes was assessed for LIANA recovery or ligand-receptor catalog membership. This separated recovery of candidate communication axes from response programs outside ligand-receptor catalog scope.

### Autocrine Potential Analysis

Chemokine and receptor expression patterns were summarized across KHYG-1 and K562 dyad populations at 2 hours. Receptor expression was used to distinguish candidate heterotypic KHYG-1-K562 axes from NK-to-NK/autocrine-like axes. Directional interaction summaries were generated from LIANA-ranked interactions and interpreted as expression-supported signaling potential rather than direct evidence of signaling flux or pathway causality.

### Foundation-Model Reconstruction Analysis

Geneformer, scGPT, and scFoundation were evaluated by reconstruction analysis using in-experiment singlets as matched reference cells and Nanovial dyads as held-out contact-associated states. Reconstruction errors were compared across singlet reference cells, 0-hour Nanovial dyads, and 2-hour Nanovial dyads. Group differences were evaluated using Mann-Whitney tests, and activated-dyad penalties were summarized relative to singlet reference-cell reconstruction error.

## Data Availability

Processed data and an interactive view of the results are available through Partillion Bioscience at https://www.partillion.com/datasets. The Cell-Cell-Seq Data Playground (https://www.partillion.com/datasets/cell-cell-seq-nk-tumor-dyads) provides an interactive resource for exploring the full result space associated with this manuscript, including differential expression, cross-cell gene-gene correlations, and comparisons with inferred signaling analyses.

## Author Contributions

J.L. and J.d.R. conceived the study and developed the Cell-Cell-Seq workflow. J.L. and D.S. performed Nanovial loading, co-culture, flow cytometry, sorting, imaging, and single-cell library experiments. J.L. performed single-cell data processing, annotation, ligand-receptor benchmarking, and foundation-model reconstruction analysis; J.L. and D.S. performed differential-expression analysis. J.L. generated initial data plots, and J.d.R. prepared the final figures and illustrations. J.L. and J.d.R. wrote the manuscript with input from all authors. D.D.C. and J.d.R. supervised the work.

