## Supplementary Information for "Cell-Cell-Seq resolves contact-associated NK cell activation in defined tumor cell dyads"

### Supplementary Figures

**Supplementary Figure 1.** Nanovials constrain Jurkat-Raji interactions in a bispecific T-cell engager co-culture model.

**Supplementary Figure 2.** Sequencing quality-control metrics for suspension co-culture, Nanovial singlets, and Nanovial dyads.

**Supplementary Figure 3.** Orthogonal annotation and demultiplexing support Cell-Cell-Seq replicate 2 dyad assignments.

**Supplementary Figure 4.** Population maps and lineage scores for Nanovial and suspension co-culture samples.

**Supplementary Figure 5.** Replicate conservation of the Cell-Cell-Seq activation response.

### Supplementary Videos

**Supplementary Video 1.** Time-lapse imaging of NK–K562 suspension (plate) co-culture, full field.

**Supplementary Video 2.** Time-lapse imaging of NK–K562 suspension (plate) co-culture, cropped view.

**Supplementary Video 3.** Time-lapse imaging of NK–K562 Nanovial co-culture, full field.

**Supplementary Video 4.** Time-lapse imaging of NK–K562 Nanovial co-culture, cropped view.

**Supplementary Video 5.** Time-lapse imaging of Jurkat–Raji co-culture in suspension with a bispecific T-cell engager (blinatumomab biosimilar).

**Supplementary Video 6.** Time-lapse imaging of Jurkat–Raji co-culture in Nanovials with a bispecific T-cell engager.

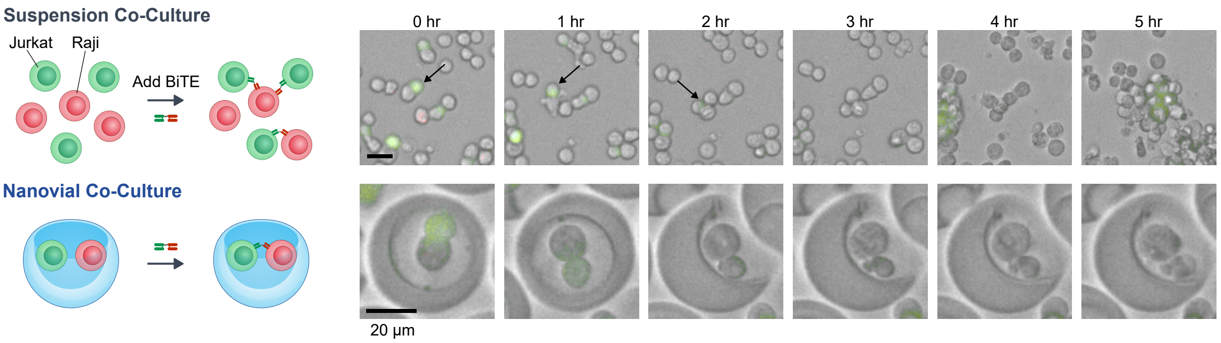


**Supplementary Figure 1. Nanovials constrain Jurkat-Raji interactions in a bispecific T-cell engager co-culture model.** Representative time-lapse imaging of Jurkat-Raji co-cultures in suspension and Nanovials after addition of a blinatumomab biosimilar. Suspension co-culture shows changing cell neighborhoods and aggregate formation over the imaging window, whereas Nanovial co-culture maintains isolated cell pairs in a local cavity. This experiment extends the physical-control argument for Nanovial co-culture beyond the KHYG-1-K562 model.

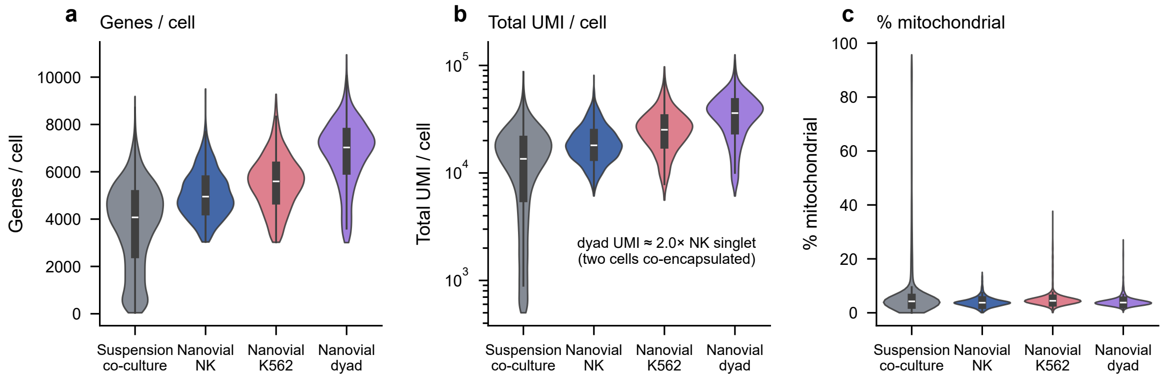


**Supplementary Figure 2. Sequencing quality-control metrics for suspension co-culture, Nanovial singlets, and Nanovial dyads. a,** Detected genes per cell, **b,** total UMI per cell, and **c,** mitochondrial read fraction, shown as violin plots across suspension co-culture, Nanovial NK singlets, Nanovial K562 singlets, and Nanovial dyads. Nanovial dyads show increased transcript recovery relative to singlets (total UMI ≈ 2× Nanovial NK singlets), consistent with two cells contributing RNA to the same droplet barcode, while mitochondrial fractions remain within the expected range after quality filtering.

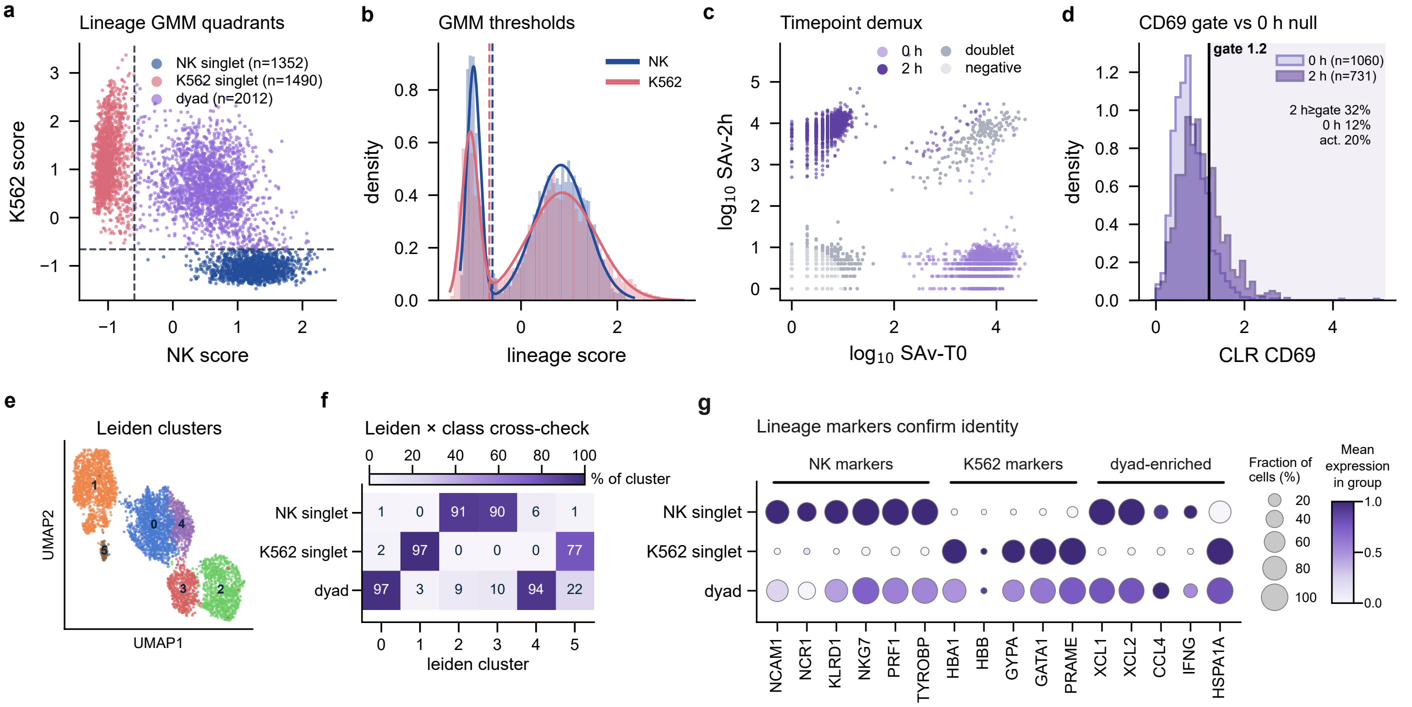


**Supplementary Figure 3. Orthogonal annotation and demultiplexing support Cell-Cell-Seq replicate 2 dyad assignments. a,** Gaussian-mixture-model lineage quadrants based on NK and K562 lineage scores assign NK singlets, K562 singlets, and dyad-associated events. **b,** Distribution of lineage scores and corresponding GMM thresholds for NK and K562 assignment. **c,** Oligo-streptavidin sample-tag demultiplexing separates 0 h, 2 h, doublet, and negative events. **d,** CD69 CLR distribution in 0 h and 2 h dyads, showing the activation gate used for cross-experiment comparisons. **e,** Leiden clusters in the pooled expression space. **f,** Cross-check of Leiden clusters against lineage classes. **g,** Dot plot of NK, K562, and dyad-enriched marker genes, confirming lineage assignments and mixed dyad identity.

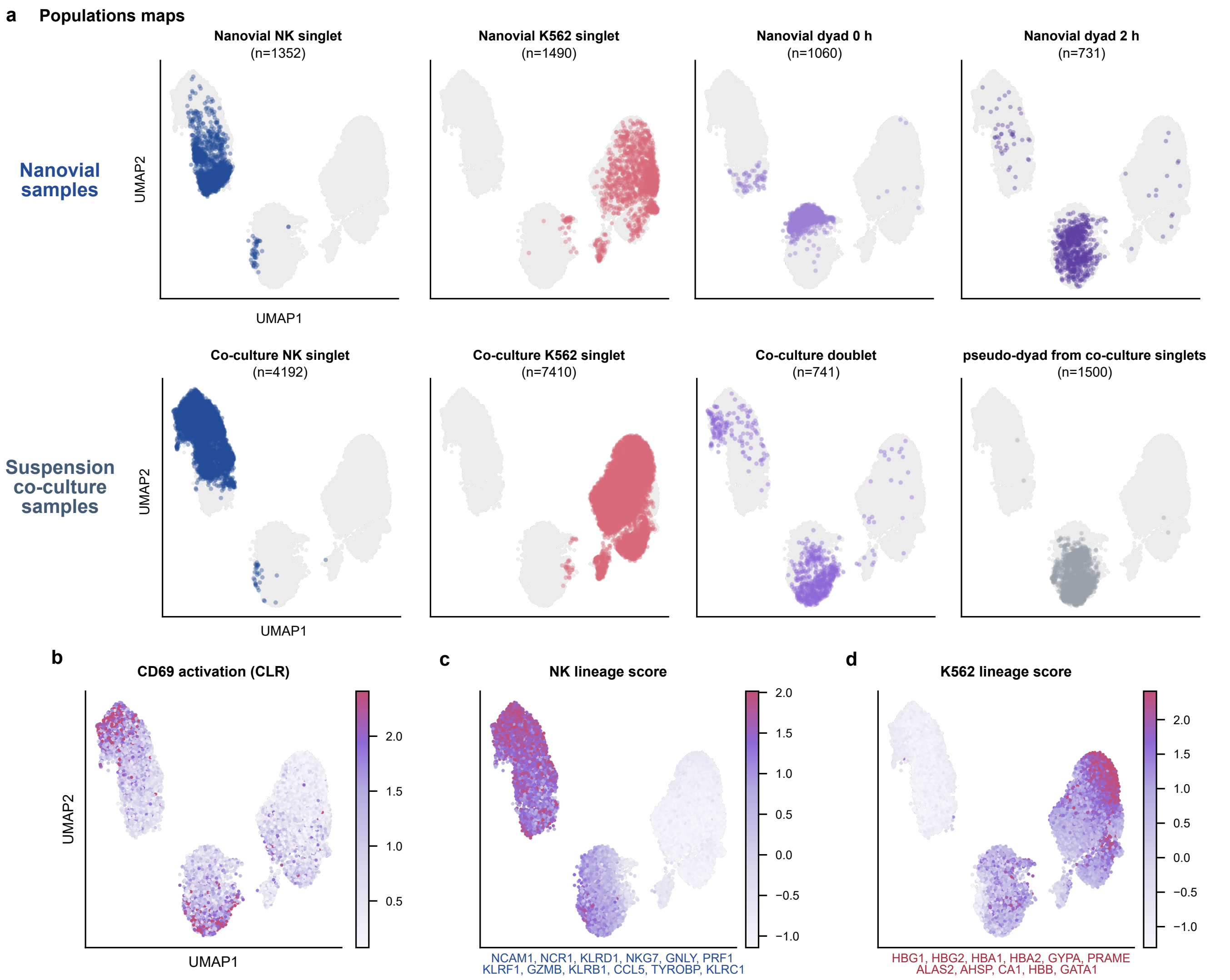


**Supplementary Figure 4. Population maps and lineage scores for Nanovial and suspension co-culture samples. a,** UMAP population maps: Nanovial NK singlets, K562 singlets, and 0 h and 2 h Nanovial dyads (top row), and suspension co-culture NK singlets, K562 singlets, doublets, and pseudo-dyads generated from co-culture singlets (bottom row). **b,** CD69 feature-barcode activation signal (CLR) across the integrated map. **c,** NK lineage-score map. **d,** K562 lineage-score map. Nanovial dyads and suspension doublets retain mixed NK/K562 lineage signal, whereas singlet populations occupy lineage-specific regions.

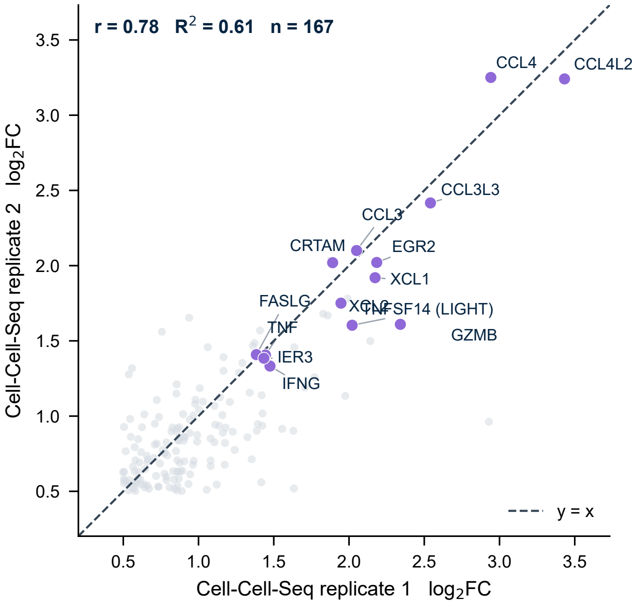


**Supplementary Figure 5. Replicate conservation of the Cell-Cell-Seq activation response.** Scatter plot comparing activation-associated log2 fold changes between Cell-Cell-Seq replicate 1 and replicate 2. Genes significant in both replicates show concordant effect sizes, including chemokines, cytotoxicity genes, immediate-early response genes, and TNFSF14/LIGHT. The conserved response supports reproducibility of the KHYG-1-K562 dyad activation program across independent Cell-Cell-Seq experiments.
